# Genome-scale detection of positive selection in 9 primates predicts human-virus evolutionary conflicts

**DOI:** 10.1101/131680

**Authors:** Robin van der Lee, Laurens Wiel, Teunis J.P. van Dam, Martijn A. Huynen

## Abstract

Hotspots of rapid genome evolution hold clues about human adaptation. Here, we present a comparative analysis of nine whole-genome sequenced primates to identify high-confidence targets of positive selection. We find strong statistical evidence for positive selection acting on 331 protein-coding genes (3%), pinpointing 934 adaptively evolving codons (0.014%). Our stringent procedure and quality control of alignments and evolutionary inferences reveal substantial artefacts (20% of initial predictions) that have inflated previous estimates of positive selection, the large majority relating to transcript definitions (61%) or gene models (38%). Our final set of 331 positively selected genes (PSG) are strongly enriched for innate and adaptive immune functions, secreted and cell membrane proteins (e.g. pattern recognition, complement, cytokine pathways, defensins, immune receptors, MHC, Siglecs). We also find evidence for positive selection in reproduction, chromosome segregation and meiosis (e.g. centromere-associated *CENPO, CENPT*), apolipoproteins, smell/taste receptors, and proteins interacting with mitochondrial-encoded molecules. Focusing on the virus-host interaction, we retrieve most evolutionary conflicts known to influence antiviral activity (e.g. *TRIM5*, *MAVS*, *SAMHD1*, tetherin) and predict 70 novel cases through integration with virus-host interaction data (virus-human PPIs, immune cell expression, infection screens). Protein structure analysis identifies positive selection in the interaction interfaces between viruses and their human cellular receptors (*CD4* – HIV; *CD46* [MCP] – measles, adenoviruses; *CD55* [DAF] – picornaviruses). Finally, the primate PSG consistently show high sequence variation in human exomes, suggesting ongoing evolution. Our curated dataset of positively selected genes and positions, available at http://www.cmbi.umcn.nl/∼rvdlee/positive_selection/, is a rich source for studying the genetics underlying human (antiviral) phenotypes.

## Introduction

Conservation of structure and sequence often indicate biological function. Rapidly evolving sequence features may however also indicate function, as they may reveal molecular adaptations to new selection pressures. But what drives these rapid genetic changes during evolution? Can these changes explain the specific phenotypes of species or individuals, such as a differential susceptibility to viruses?

Immunity genes contain the strongest signatures of rapid evolution due to positive Darwinian selection (Vallender and Lahn 2004; Bustamante et al. 2005; Chimpanzee Sequencing and Analysis Consortium 2005; Nielsen et al. 2005; Voight et al. 2006; Rhesus Macaque Genome Sequencing and Analysis Consortium et al. 2007; Kosiol et al. 2008; George et al. 2011; Fu and Akey 2013; van der Lee et al. 2015; Deschamps et al. 2016; Enard et al. 2016). Pathogens continuously invent new ways to evade, counteract and suppress the immune response of their hosts, thereby acting as major drivers of the observed adaptive evolution of immune systems (Holmes 2004; Daugherty and Malik 2012). Numerous proteins involved in the virus-host interaction have been demonstrated to be in genetic conflict with their interacting viral proteins, a phenomenon that has been likened to a virus-host ‘arms race’ (Daugherty and Malik 2012). Such studies have generally focused on a single gene or gene family of interest sequenced across a large number of species. Evolutionary analyses can then predict which genes and codons may be involved in virus interactions. For example, mitochondrial antiviral signaling protein (MAVS) is a central signaling hub in the RIG-I-like receptor (RLR) pathway, which recognizes infections of a wide range of viruses from the presence of their RNA in the cytosol. Analysis of the MAVS gene in 21 primates identified several positions that have evolved under recurrent strong positive selection and turned out to be critical for resisting cleavage by Hepatitis C virus (Patel et al. 2012). Other examples of immunity genes showing evolutionary divergence that directly impacts the ability to restrict viral replication include TRIM5α (Sawyer et al. 2005), PKR (Elde et al. 2009) and MxA (Mitchell et al. 2012).

Positive selection can be detected through comparative analysis of protein-coding DNA sequences from multiple species (Yang and Bielawski 2000; Fu and Akey 2013). Markov models of codon evolution combined with maximum likelihood (ML) methods (Yang 2007) can analyze alignments of orthologous sequences to identify genes, codons and lineages that show an excess of nonsynonymous substitutions (mutations in the DNA that cause changes to the protein) compared to synonymous (‘silent’) substitutions (*d*_N_/*d*_S_ ratio or ω, see **Text S1** for a detailed explanation). Successful application requires many steps (Yang et al. 2000; Daugherty and Malik 2012)(**Text S1**), including: (i) the identification of orthologous sequences, sampled from species across an appropriate evolutionary distance (distant enough to show variation, but not too divergent to prevent saturation); (ii) accurate alignment and phylogenetic tree reconstruction; (iii) parameterization of the ML model. While studies of positive selection on individual genes have achieved reliable results, estimates of positive selection in whole genomes have been substantially affected by unreliabilities in sequencing, gene models, annotation and misalignment (Wong et al. 2008; Schneider et al. 2009; Fletcher and Yang 2010; Markova-Raina and Petrov 2011; Jordan and Goldman 2012; Privman et al. 2012; Moretti et al. 2014). In addition to the analysis of genomes from different species, the increasing availability of genomes from human individuals and populations provides new opportunities to systematically analyze more recent sequence variation (Fu and Akey 2013; Lek et al. 2016).

In this study, we performed comparative evolutionary analyses of recent whole-genome sequenced primates as well as of human genetic variation to identify high-confidence targets of positive selection. Our findings provide insights into the biological systems that have undergone molecular adaptation in primate evolution. Interrogation of the positively selected genes with structural and genomic data describing virus-host interactions provides insights into potential determinants of viral infection and predicts new virus-human evolutionary conflicts.

## Results

To obtain a confident dataset of positively selected genes relevant to human biology and infectious disease, we investigated protein-coding DNA sequences from nine simian (‘higher’) primates for which whole-genome sequences are available (**Table S1**, five genomes released in 2012 or later (Rogers and Gibbs 2014)). This set consists of five hominoids (‘apes’; human, chimpanzee, gorilla, orangutan, gibbon), three Old World Monkeys (macaque, baboon, vervet) and one New World monkey (marmoset), spanning an estimated 36–50 million years of evolutionary divergence (Perelman et al. 2011).

### A reliable procedure for conservative inference of positive selection

Given the high incidence of false positives in large-scale detection of positive selection reported in literature (Wong et al. 2008; Schneider et al. 2009; Fletcher and Yang 2010; Markova-Raina and Petrov 2011; Jordan and Goldman 2012; Privman et al. 2012; Moretti et al. 2014), we developed a stringent six-stage procedure that we subjected to rigorous manual curation and quality control at all steps. This section and Figure 1 summarize the procedure. More details are available in the **Methods** section.

**Figure 1.**
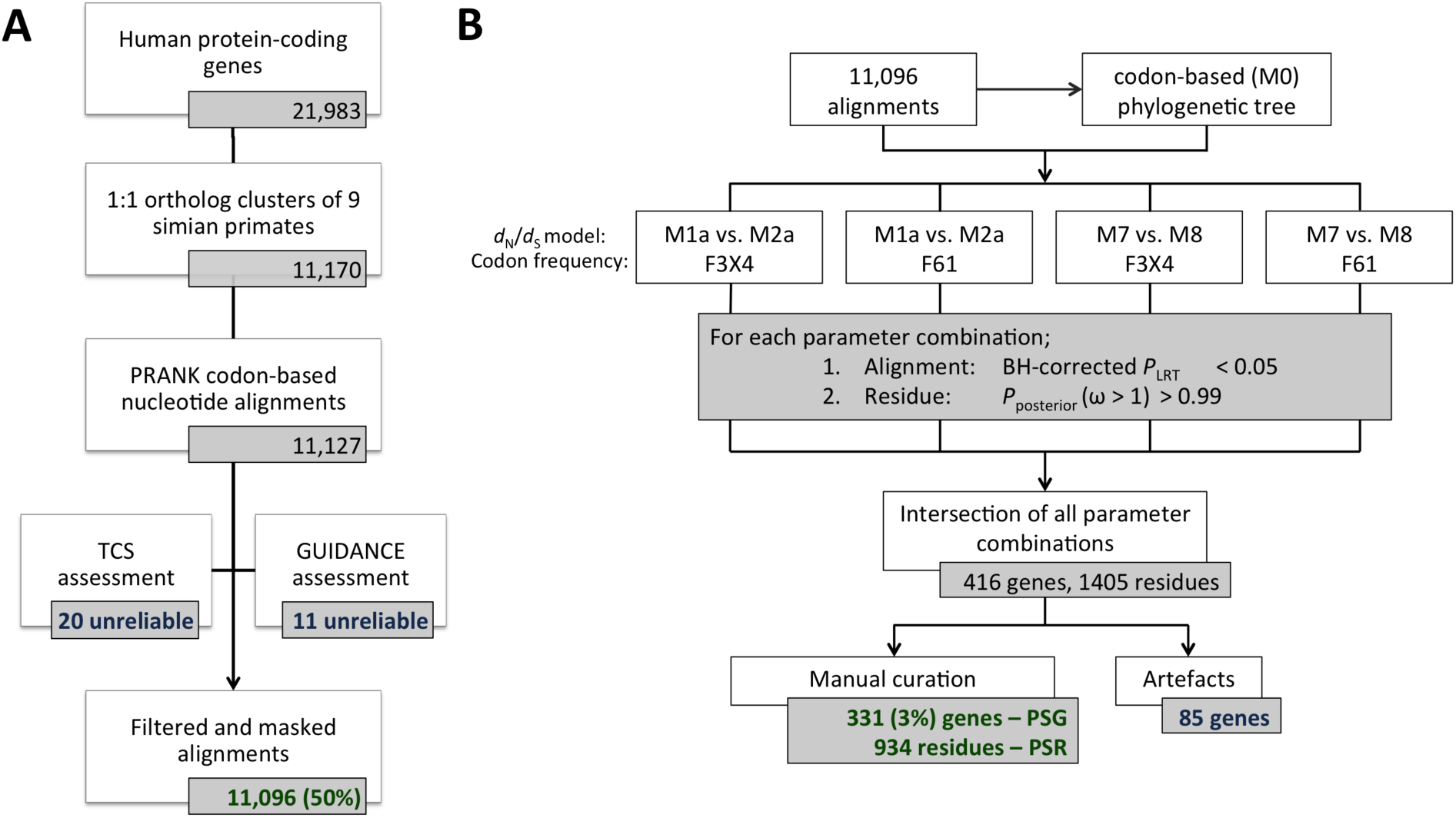
Large-scale comparative evolutionary analysis procedure for conservative inference of positively selected genes and codons. (**A**) Orthology inference, multiple sequence alignment, and alignment filtering/masking steps. (**B**) Maximum likelihood evolutionary analyses, statistics, and manual curation steps. Both stages were subject to rigorous curation and quality control at all steps. See main text and **Methods** for details. See **Figure S2** for detailed results of the various ML parameter combinations (B). See **Figure 3** for a breakdown of the artefacts (B).

To minimize artefacts, we prioritized high precision over sensitivity (i.e. potentially missing interesting sites). (i) To limit the influence of evolutionary processes other than divergence of orthologous codons (e.g. gene duplications), we only assessed clusters of one-to-one orthologous genes (**Table S2**). (ii) To reduce alignment of nonhomologous codons, a common issue leading to inflated estimates of positive selection (Schneider et al. 2009; Fletcher and Yang 2010; Markova-Raina and Petrov 2011; Jordan and Goldman 2012; Privman et al. 2012; Moretti et al. 2014), we computed multiple sequence alignments using PRANK (Löytynoja and Goldman 2008). (iii) To achieve maximum alignment quality, we masked low-confidence codons and columns, and filtered out alignments based on the GUIDANCE and TCS algorithms, two distinct concepts for assessing reliability (Penn et al. 2010; Chang et al. 2014). These steps resulted in 11,096 alignments, representing about half of the human protein-coding genes (Figure 1A, Table 1). Detailed inspection revealed that the alignments are of good overall quality, with the major improvements gained by using PRANK over other alignment methods. The masking procedures filter out most of the remaining ambiguities.

**Table 1.** Positive selection statistics.

| Description | Value | Other |
| --- | --- | --- |
| *Species investigated* | 9 | human, chimpanzee, gorilla, orangutan, gibbon, macaque, baboon, vervet, marmoset |
| *One-to-one ortholog clusters analyzed for positive selection* | 11,096 | 50% of human protein-coding genes; $6.6 \times 10^6$ codons human; $5.7 \times 10^7$ codons total |
| *Overall $d_N/d_S$ rate (across 11,096 clusters)* | 0.21 | $d_N = 0.0477$, $d_S = 0.2235$ |
| *Positively selected genes (PSG)<sup>a</sup>* | 331 | 3% of genes tested |
| *Positively selected residues (PSR)* | 934 | 0.5% of $1.9 \times 10^5$ PSG codons; 0.014% of $6.6 \times 10^6$ tested human codons |
| *PSG $d_N/d_S$ rate<sup>b</sup>* | 6.85 (median) | 2.10 (minimum) |

<sup>a</sup> Genes for which across all four evolutionary models tested: (i) the selection model gives a significantly better fit than the neutral model and (ii) that contain at least one significant PSR (**Methods**).
<sup>b</sup> Calculated from the per-PSG averages of the $d_N/d_S$ rates estimated in the four evolutionary models tested (**Methods**).

Next, we used *d*_N_/*d*_S_-based codon substitution maximum likelihood (ML) models (Yang 2007) to infer positive selection acting on genes and codons (Figure 1B, **Methods**, **Text S1**). This requires an estimation of the overall evolutionary divergence between the primate species. (iv) To best reflect this distance, we used the 11,096 one-to-one ortholog alignments to construct a single codon-based reference tree for use in all ML analyses (Figure 2, **S1**; see also next section). (v) To ensure that we studied only the strongest signatures of positive selection, we analyzed four combinations of evolutionary model parameters and required genes to test significant across all of them (*P* < 0.05, after Benjamini-Hochberg correction for testing 11,096 alignments). (vi) Finally, we obtained the set of positively selected codons (and their corresponding amino acid residues) using stringent Bayesian calculations (*P*_posterior_ > 0.99). These steps resulted in 416 apparent Positive Selected Genes (aPSG) inferred to contain at least one apparent Positively Selected Residue (aPSR; 1405 in total, **Figure S2**, **Table S3**).

**Figure 2.**
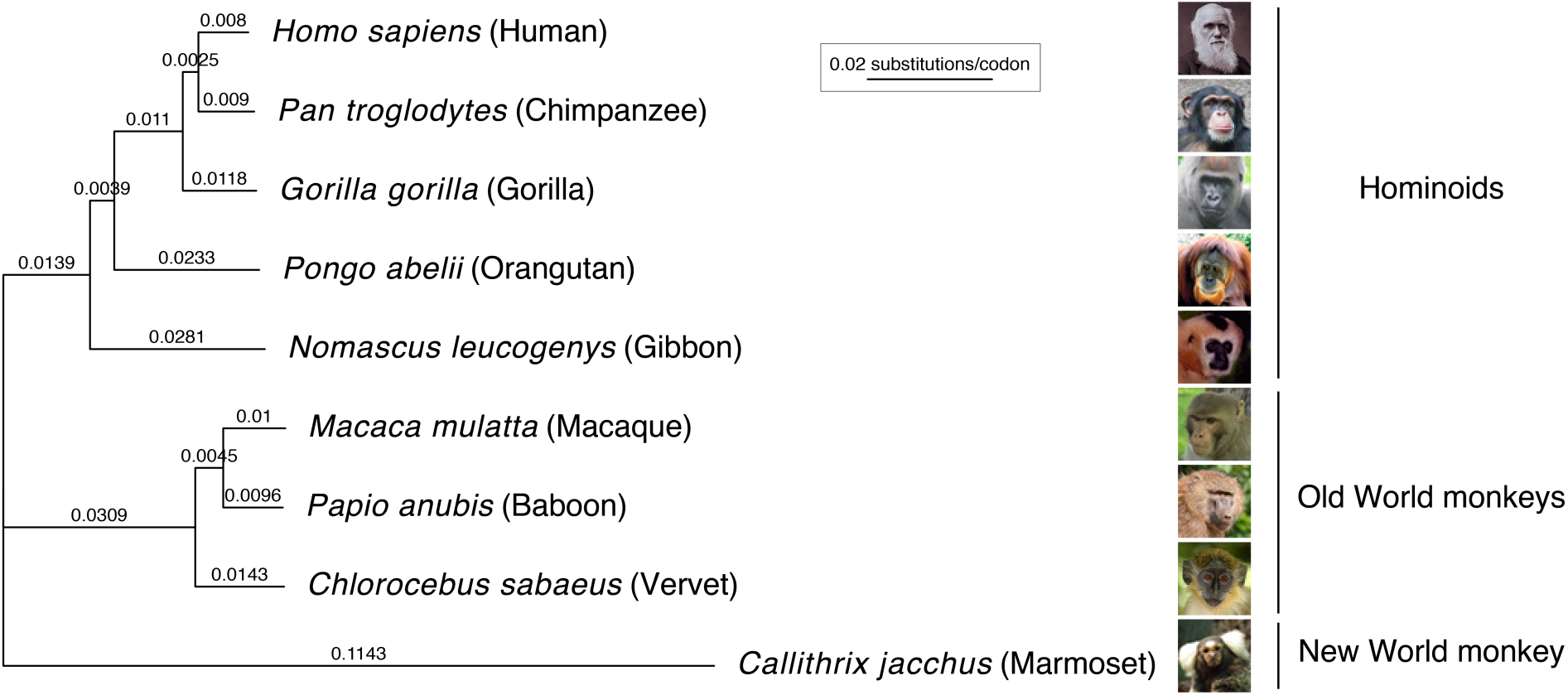
Codon-based phylogenetic tree of nine simian primates. Branch lengths (nucleotide substitutions per codon) were estimated using the codeml M0 (F61) evolutionary model on the concatenated, masked alignment of 11,096 protein-coding, one-to-one orthologous genes of the nine primates studied. See **Figure S1** for species image credits.

### Evolutionary models estimate substitution rates in primate protein-coding genes

The overall *d*_N_/*d*_S_ rate across our one-to-one protein-coding orthologs is 0.21 (*d*_N_ = 0.0477, *d*_S_ = 0.2235; Table 1), consistent with average strong purifying selection. Our primate phylogenetic tree, calculated for the ML analyses and based on all included alignments, has identical topology to the well-established primate phylogeny (Figure 2, **S1**)(Perelman et al. 2011). The estimated substitution rates per nucleotide follow the expected pattern, depending on the fraction of noncoding sequences used for tree reconstruction: the nucleotide-converted branch lengths of our coding sequence tree are a factor 0.87 shorter (median of all branches) than those of a phylogeny based on genomic regions of 54 primate genes (consisting half of noncoding segments)(Perelman et al. 2011), and a factor 0.46 shorter than a tree based on whole-genome alignment (Yates et al. 2016)(**Figure S1**). Thus, our phylogenetic tree informs on the rate of protein-coding sequence divergence observed in primates.

### Rigorous quality control reveals artefacts in 20% of apparent PSG

To assess the reliability of our procedure and estimate the impact of common errors in large-scale sequence analysis, we performed systematic manual inspection of all 1405 aPSR and 416 aPSG. Based on the results, we implemented filters for automatic detection of false predictions of positive selection (referred to as artefacts; **Methods**). Although the large majority of predictions are reliable, 85 aPSG (20%) contain artefacts.

The artefacts we encountered fall into five classes (Figure 3A, **Table S3**). (I) Orthology inference or gene clustering errors (occurring in 8 of 85 problem cases, 9%). One gene cluster consists of the *TRIM60* sequences of seven primates plus the *TRIM75* sequences of baboon and gibbon. *TRIM60* and *TRIM75* are close paralogs (46% identical amino acids, Figure 3B) encoded within a 43kb region on chromosome 4 in human. *TRIM75P* is an annotated pseudogene in human and *TRIM60P* is a predicted pseudogene in gibbon, despite both appearing like fully functional genes: they lack frame-shift mutations or premature stop codons and remain highly similar to the corresponding non-pseudogenic copy in the other species (97-98% identical amino acids). Clustering of the outparalogs *TRIM60* and *TRIM75* led to artificially high substitution rates and an abundance of apparent PSR across the full alignment (**Figure S3**). (II) Alternative transcript definitions (52 cases, 61%). In most cases this leads to alignment of divergent exons (**Figure S3**). The *CALU* gene cluster consists of the coding sequences from two alternative isoforms that include either one of two genomic neighboring and homologous exons (Figure 3C). Given their strong sequence conservation (64% identical amino acids), these mutually exclusive exons likely originated from a tandem exon duplication event (Letunic et al. 2002). For six primates the *CALU* gene model predicted a single transcript isoform that contains the first of the tandem exons, which made these sequences inconsistent with the alternative transcript selected for the other primates. (III) Unreliable N-/C-termini (32 cases, 38%). When they are sufficiently similar, alternative non-homologous translation starts/stops or alternative exon boundaries may still be aligned. Such cases appear as divergent, high *d*_N_/*d*_S_ codons (**Figure S3**). (IV) Alignment ambiguities (4 cases, 5%). These are spurious cases of short, hard-to-align sequence regions, usually surrounding residues that are masked due to low alignment quality (**Table S3**). Without independent data (e.g. more sequences, aligned 3D structures) it remains challenging to determine the correct alignment. (V) Other issues (3 cases, 4%). These are apparent PSR with high posterior probabilities in all ML model parameter combinations, but upon closer inspection are untrustworthy (**Table S3**). One site for example consisted of distinct codons distributed across the tree (6x AGC, 3x TCA), which were inferred to evolve under high *d*_N_/*d*_S_ despite all encoding serine residues.

**Figure 3.**
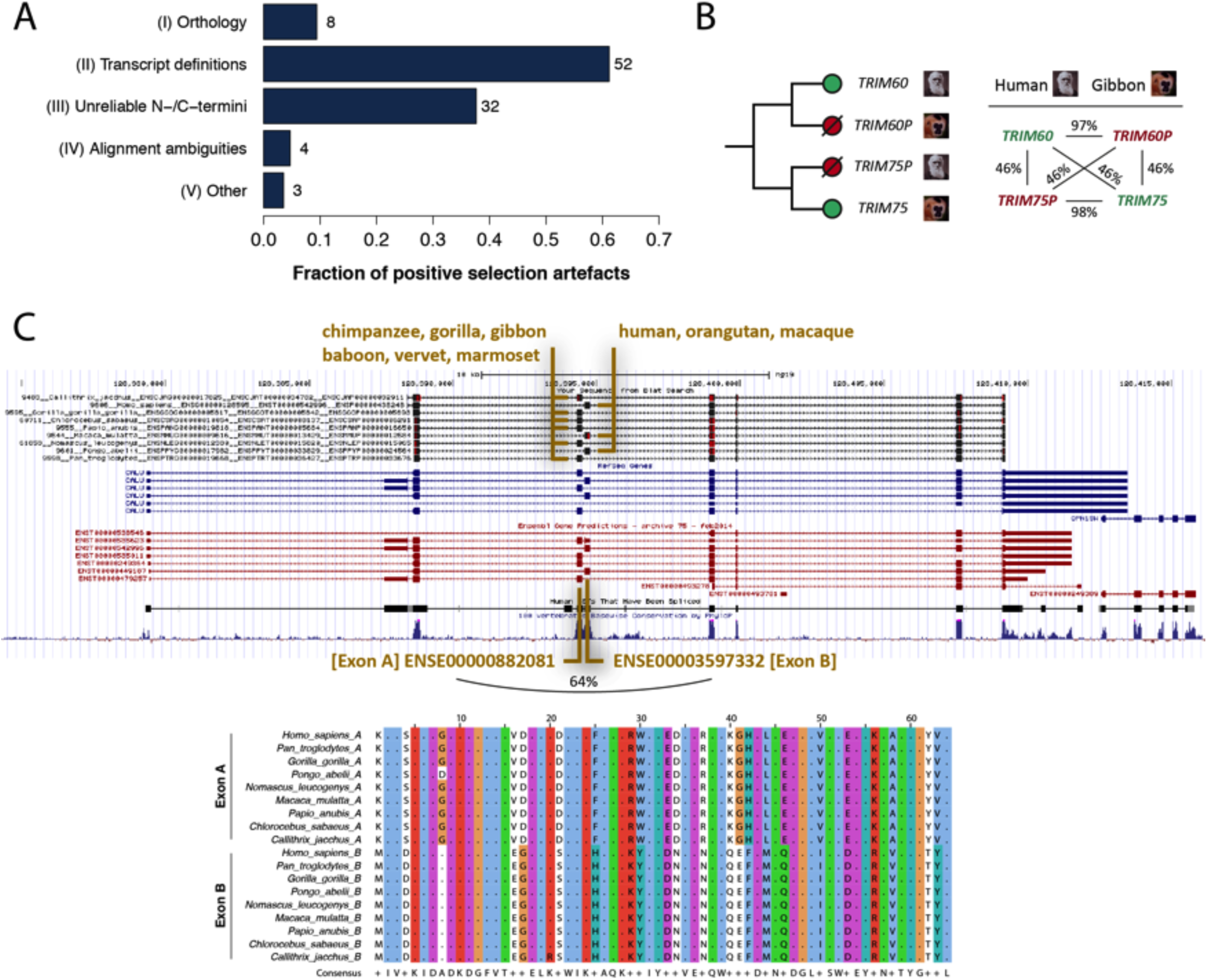
Positive selection artefacts. (**A**) Five classes of artefacts. Note that each alignment may be affected by more than one type of problem. (**B**) A type-I problem (orthology) involving the clustering of outparalogs *TRIM60* and *TRIM75*, which are highly similar within and between primates. *TRIM75P* in human and *TRIM60P* in gibbon are annotated pseudogenes. (**C**) A type-II problem (transcript definitions) involving mutually exclusive, tandem duplicated exons. [top] BLAT alignment (UCSC Genome Browser (Rosenbloom et al. 2015)) of cDNA sequences of the nine primates (black tracks) to the 45kb *CALU* genomic region on human chr7. [bottom] Multiple alignment of the translated sequences of Exons A and B. Percentages refer to pairwise identities on the protein level as determined by global alignment. See **Figure S3** for details and other examples.

### GC-biased gene conversion does not systematically affect the PSG

GC-biased gene conversion (gBGC) may be an alternative explanation for the accelerated evolution of some PSG (Galtier and Duret 2007). gBGC leads to increased GC content in meiotic recombination hotspots, which may inflate *d*_N_/*d*_S_ estimates (Ratnakumar et al. 2010). Genes affected by gBGC are expected to have substitution patterns more biased towards GC than genes evolving under positive selection, particularly in the selectively neutral fourfold degenerate (FFD) sites.

We analyzed our PSG to determine whether they are affected by gBGC. First, we found that PSG have significantly lower rather than higher GC contents compared to non-PSG in all studied primates, both across the full coding sequence as well as in FFD sites (human: median GC_FFD_ = 0.518 (PSG), 0.578 (non-PSG), *P* = 5.3 × 10^−6^, Mann-Whitney *U* test; **Figure S4**). Second, amino acids enriched for PSR positions correlate only marginally with GC content (Pearson’s *R* = 0.14, *P* = 0.56; **Figure S5**). Third, PSG overlap only marginally with human recombination hotspots (7-8% for both PSG and non-PSG), as well as with genomic regions predicted to be affected by gBGC (Capra et al. 2013)(9% of PSG vs. 11% of non-PSG). Moreover, our site-specific PSG inferences are based on substitution patterns across a primate phylogeny covering at least 36 million years of evolution, which far outdates the rapid turnover rate of recombination hotspots (e.g. even human and chimp hotspot rarely overlap (Winckler et al. 2005)), strongly reducing the influence of gBGC (Ratnakumar et al. 2010). Together these data suggest that although gBGC may affect individual PSG, the adaptive signatures in the large majority of PSG are not caused by gBGC but are likely the result of positive selection.

### Strong statistical evidence for positive selection in 3% of human protein-coding genes

Removal of the 85 detected artefacts resulted in a final, curated set of 331 human genes with extensive statistical evidence for positive selection across nine primates (331 PSG, 3% of 11,096 ortholog clusters analyzed; Table 1, **S4**). The PSG are distributed evenly across the human genome (**Figure S6**). They have a median *d*_N_/*d*_S_ rate of 6.85 (minimum 2.10), consistent with the strong positive selection observed across multiple evolutionary models. 106 of the 331 PSG (32%) had been reported in an overview of previous genome-wide studies for positive selection, which include comparisons of different species as well as studies of human variation (Fu and Akey 2013). The 331 PSG contain 934 positively selected residues/codons (PSR). The 934 PSR comprise 0.014% of all tested human codons and 0.5% of all PSG codons, with an average of 2.8 PSR per PSG (Table 1, **S5**). Over half (53%) of PSG have a single PSR and 14 genes have 12 or more PSR with a maximum of 38: from high to low, *MUC13, PASD1, NAPSA, PTPRC, APOL6, MS4A12, CD59, SCGB1D2, PIP, CFH, RARRES3, OAS1, TSPAN8, TRIM5*. We observed a notable enrichment of arginine and histidine among the 934 PSR (**Figure S7**), which may indicate protein functionality through charged interactions (Tsai et al. 1997).

### Immune pathways and functions are abundant among PSG

To gain more insight into the processes that evolved under positive selective pressure in primates, we investigated our 331 PSG for molecular functions, biological pathways and other properties. PSG are strongly enriched for a variety of immune-related pathways and gene ontology categories of both innate and adaptive nature, including functions in: inflammation, complement cascades, hematopoiesis, B-and T-cell immunity and the defense response against bacteria and viruses (**Table S6**, **S7**). Overlap analysis confirmed the enrichment of innate immune functions among PSG (49/331 genes [15%], 2.8-fold enriched compared to non-PSG, *P* = 2.8 × 10^−10^, two-tailed Fisher’s exact test) and pattern recognition pathway components (12 genes, **Table S8**). These include *TLR1, TLR8, MAVS, IFI16, CASP10, TRIM5, OAS1, RNASEL, PGLYRP1, NLRP11, CLEC1A* and *CLEC4A* (Table 2). In addition, many PSG encode transmembrane (65 genes [20%], 1.4-fold, *P* = 6.4 × 10^−3^) or secreted proteins (73 genes [22%], 2.5-fold, *P* = 1.4 × 10^−12^), as is also indicated by the abundance of enriched terms associated with: extracellular and cell surface localization, receptor activity, signal peptide, disulfide bond and glycosylation (**Table S6**, **S7**).

**Table 2.**
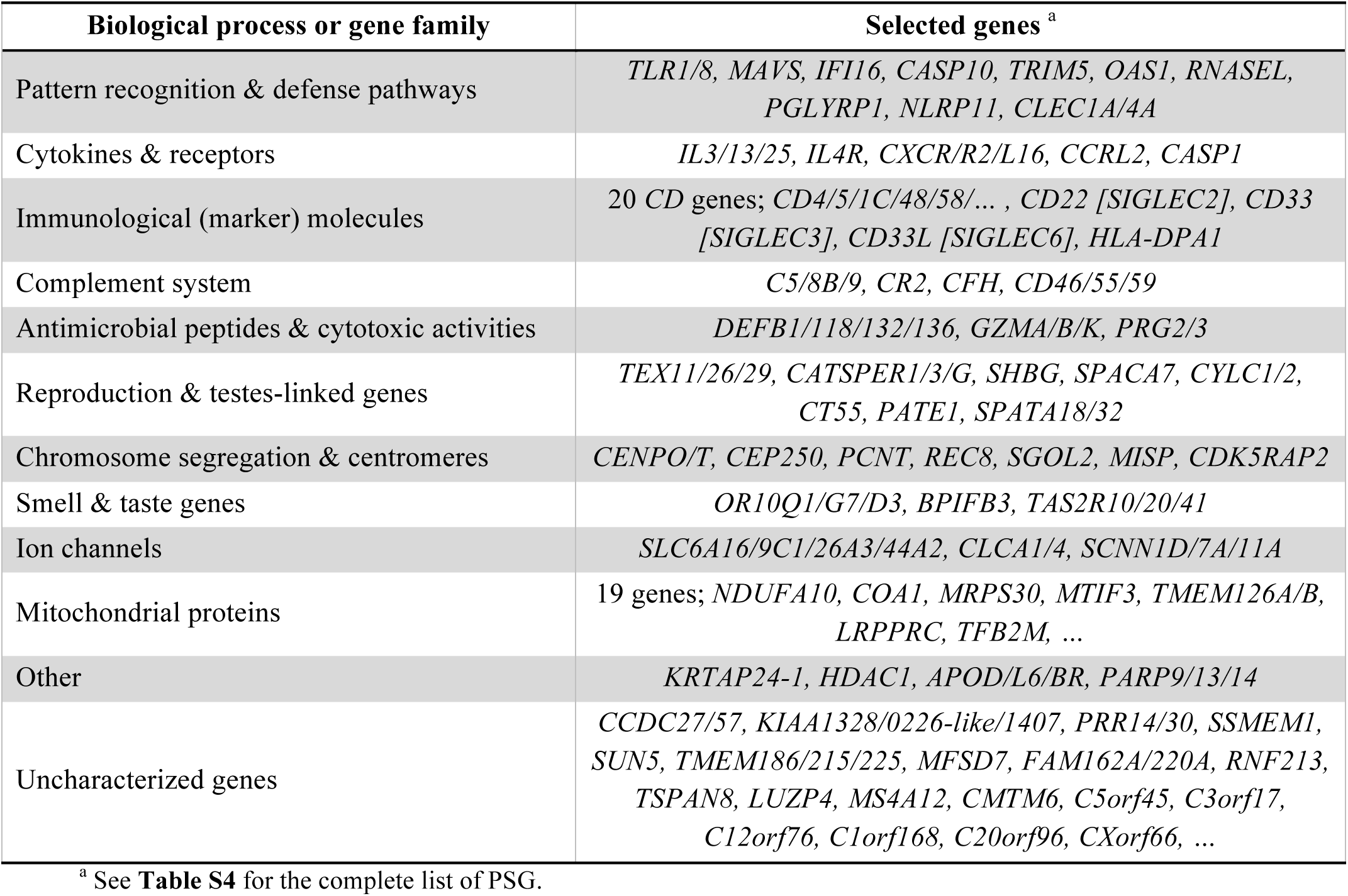
Positively selected processes and gene families.

Detailed examination revealed a range of other noteworthy genes and processes among the PSG. These include cytokines and their receptors (e.g. *IL3, IL4R* and *CASP1*, which activates IL-1β and IL-18), various immunological marker molecules (20 ‘cluster of differentiation’ genes, including *CD4/5/48* and the sialic acid binding Ig-like lectins *SIGLEC2/3/6*), an MHC class II subunit (*HLA-DPA1*), genes with antimicrobial activity (defensins, granzymes), olfactory and taste receptors (*IR10Q1, TAS2R20*), ion channels (solute carrier, chloride, sodium channel families), a keratin associated protein (*KRTAP24-1*), poly (ADP-ribose) polymerases (*PARP9/13/14*), apolipoproteins (*APOD/L6/B*), and various genes of unknown function (∼20-25% of 331 PSG; Table 2, **S8**). The PSG further contain 19 nuclear genes encoding mitochondrial proteins (e.g. OXPHOS complex I subunit *NDUFA10* and assembly factor *TMEM126B*), potentially indicating compensatory evolution between nuclear and mitochondrial genomes (**Discussion**). Interestingly, we also found considerable evidence for positive selection associated with reproduction: genes linked to spermatogenesis and testes (e.g. *TEX11, CATSPER1, SPATA32*), and genes involved in the chromatin structure of the centromere, the kinetochore, chromosome segregation and meiosis (e.g. *CENPO/T, REC8, PCNT*; **Discussion**).

### Positive selection identifies known and novel virus-host genetic conflicts

The strong signal for immunity among the positive selection data suggests that at least some rapidly evolving sites are in a genetic conflict with one or more pathogens. To further assess the ability of the PSG data to detect such conflicts, we investigated our dataset for known virus-human evolutionary interactions. We evaluated five cases in which positively selected codons, identified through evolutionary analysis, were experimentally shown to be important for restricting viral infection. Our large-scale approach correctly identified four out of five genes (*TRIM5*, *MAVS*, *BST2* [tetherin], *SAMHD1* (Sawyer et al. 2005; Gupta et al. 2009; Laguette et al. 2012; Patel et al. 2012); MxA was not detected (Mitchell et al. 2012); **Table S9**). Moreover, despite using sequences from substantially less species (those case studies sequenced one gene in ∼20-30 primates), we retrieved many of the previously reported codons (7 previously reported out of the 12 detected by our screen for *TRIM5*, 1/1 for *MAVS*, 0/1 for *BST2*, 1/3 for *SAMHD1*). In addition to such case studies with both statistical and experimental support for antiviral positive selection, our large-scale approach also recovered PSG for which only statistical evidence for positive selection was previously reported in small-scale studies (*IFI16*, *OAS1*, *TLR8*, *PARP9/13/14*, *APOL6* (Smith and Malik 2009; Wlasiuk and Nachman 2010; Cagliani et al. 2014; Daugherty et al. 2014; Hancks et al. 2015); **Table S9**).

Next, we probed our evolutionary data for novel cases of virus-human genetic conflicts. To prioritize the PSR and PSG, we integrated them with a multitude of orthogonal datasets describing various aspects of antiviral immunity and virus-host interactions (**Table S8**), including virus-human protein interactions (Durmuş Tekir et al. 2013), functional screens of virus infection (Schmidt et al. 2013), gene expression in immune cell subtypes (Shay and Kang 2013) and virus-infected cells (van der Lee et al. 2015), and maps of recent human adaptation (Grossman et al. 2013). We found that PSG are enriched for genes showing differential expression upon infection with respiratory viruses (12 genes [4%], 3.3-fold, *P* = 5.4 × 10^−4^; **Table S8**). Many PSG are also involved in virus-human PPIs (70 genes [21%]; though not more often than non-PSG). Among the 70 PSG whose protein products interact with viruses, 49 are expressed in several or all of 14 profiled immune cell subtypes (including B-, T-, NK-cells, DCs and monocytes). 11 of these 49 also alter the course of cellular infection upon knockdown (e.g. with Hepatitis C virus [HCV], Sendai virus [SeV] or human papilloma virus 16 [HPV16]; **Table S8**). Besides containing well-described viral interactors such as *MAVS, SAMHD1* and *CD55*, those 11 PSG include poorly characterized genes that may represent novel candidate virus-host interactors. For example, *FBXO22* encodes an F-box family protein, which may be involved in ubiquitin-mediated protein degradation. *FBXO22* has not been linked to (viral) infections, other than a role in macrophage NFκB activation during *Salmonella* infection (Pilar et al. 2012). Our evolutionary data together with the virus-host interaction data suggest a role for *FBXO22* in primate HPV infections, possibly mediated by the positively selected third codon (Pro in human, chimpanzee, baboon, marmoset – Thr, Ser, Ala or Leu in gorilla, orangutan, gibbon, macaque, vervet). Ribosomal protein S29 (*RPS29*) is another example of a poorly annotated PSG. It is not only a strong target of positive selection in our study but it also localizes to a genomic locus that underwent recent human adaptation (Grossman et al. 2013). *RPS29* is expressed in 14 immune cell subsets, interacts with six different Influenza proteins, and affects HCV and SeV replication. Thus, data-driven prioritization of genes under positive selection may reveal novel cases of virus-host interactions shaped by evolutionary conflicts.

### PSR are at the structural interface between viruses and their cellular receptors

To gain further insights into the role of positive selection in viral infection, we analyzed the involvement of PSR in structurally characterized virus-host interactions. Among the PSG with strong adaptive signatures, we identified three genes that function as virus receptors and for which structures of human-virus protein complexes have been solved: *CD4*, *CD46* and *CD55*. *CD4* interacts with MHC class II molecules and is the classical surface marker of T helper cells (CD4^+^ T cells). HIV exploits CD4 as a receptor for entering host T-cells (Zhou et al. 2007). The two residues identified as positively selected by our approach, Asn77 and Ala80, are part of the CD4 V-set Ig-like domain. Both lie close to the interaction interface between CD4 and HIV envelop protein gp120, with CD4 Asn77 making extensive contact and hydrogen bonding to gp120 Ser365 (Figure 4A).

**Figure 4.**
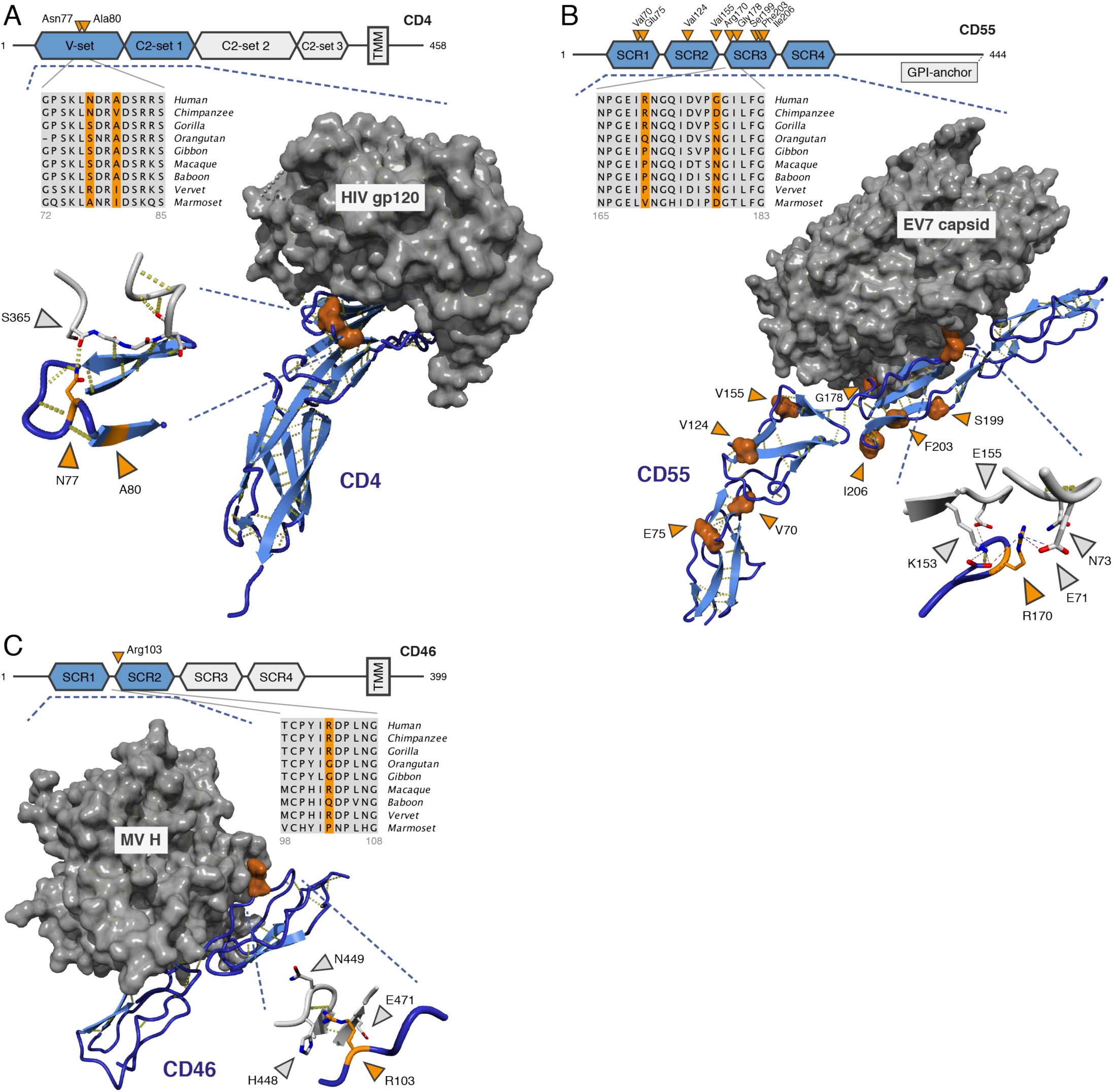
Positively selected positions in the interaction between viruses and their cellular receptors. (**A**) CD4 – HIV-1 envelop protein gp120 (2NXY (Zhou et al. 2007)). (**B**) CD55 [DAF] – echovirus 7 capsid (3IYP (Plevka et al. 2010)). (**C**) CD46 [MCP] – measles virus H (3INB (Santiago et al. 2010)). (A,C) structures of interacting proteins crystallized together as complexes; (B) EM reconstruction fitted with the individual crystal structures. Hydrogen bonds are in yellow; charge interactions are shown as dashed lines (inset of B). Blue: human receptors, orange: PSR, gray: viral proteins. Protein domains that are present in the structures are in blue and marked by the dashed lines. PSR numbering is based on the human Ensembl sequence. Viral positions correspond to sequences represented in the structures.

*CD55* [DAF] and *CD46* [MCP] are members of the regulator of complement activation (RCA) gene family that control activation of the complement cascade (21% amino acid identity; both composed of four SCR domains). *CD55* acts as an entry receptor for some picornaviruses, including several types of coxsackie-, entero-and echovirus (Bhella et al. 2004; Plevka et al. 2010). *CD55* SNPs are associated with geographical pathogen richness and *CD55* in human likely evolved under balancing selection, which maintains allelic diversity in a population (Fumagalli et al. 2009). We identified nine positively selected positions in CD55 (two in SCR domain 1, two in SCR2, five in SCR3). Arg170 and Gly178 (part of SCR3) are involved in the interaction with echoviruses (EV) 7 and 12, with Arg170 buried deep into the EV7 viral capsid (Figure 4B). A previous study reported additional interactions between picornaviruses and our CD55 PSR, including Val155 (EV7), Val124 and Ile206 (coxsackievirus B3)(Plevka et al. 2010).

*CD46* is a receptor for measles virus (MV), human herpesvirus 6 and several adenovirus subspecies (Av) (Cupelli et al. 2010; Santiago et al. 2010). Protein structure analysis revealed that the single PSR in CD46, Arg103 (SCR domain 2), makes extensive contacts with both MV hemagglutinin (His448, Asn449, Glu471; Figure 4C) and the Av type 21 fiber knob (Asn304, Ile305). Taken together, examination of the PSG with known structures suggests that positions evolving under strong positive selection may be central to the interactions between virus particles and their human cellular receptors.

### Primate PSG show ongoing change within the human population

The constantly changing spectrum of viruses and other pathogens causes a recurrent selective pressure on the human genes that interact with them (Daugherty and Malik 2012). We therefore expected many genes that evolved under positive selection in primates (i.e. our PSG) to still be undergoing adaptive evolution in recent and current human evolution. To investigate this, we analyzed patterns of human variation in the Exome Aggregation Consortium (ExAC) dataset of 60,706 exomes (Lek et al. 2016).

PSG, compared to genes that lacked between-primate signatures of positive selection (i.e. non-PSG), contain significantly more unique missense variants (*P* = 5.8 × 10^−13^), but similar amounts of unique synonymous variants (*P* = 0.23, Figure 5A). Indeed, a *d*_N_/*d*_S_-based measure of human variation, corrected for all possible missense and synonymous variants based on the codon table (**Methods**), shows that PSG have higher human *d*_N_/*d*_S_ ratios than non-PSG (median: 0.71 vs. 0.63, *P* = 1.5 × 10^−15^; Figure 5A). Similar patterns emerge when analyzing human variation at the level of allele frequencies rather than at the level of unique reported variants: PSG again show more total missense variation than non-PSG, but similar amounts of total synonymous variation (Figure 5B). PSG also contain far more ‘functional variation’ (missense, nonsense and splice variants) according to the Residual Variation Intolerance Score, which for each gene measures the deviation from the expected amount of functional variation given the background of synonymous variation present that gene (Petrovski et al. 2013)(median genome-wide RVIS percentile: 71% [PSG] vs. 45% [non-PSG], Figure 5C).

**Figure 5.**
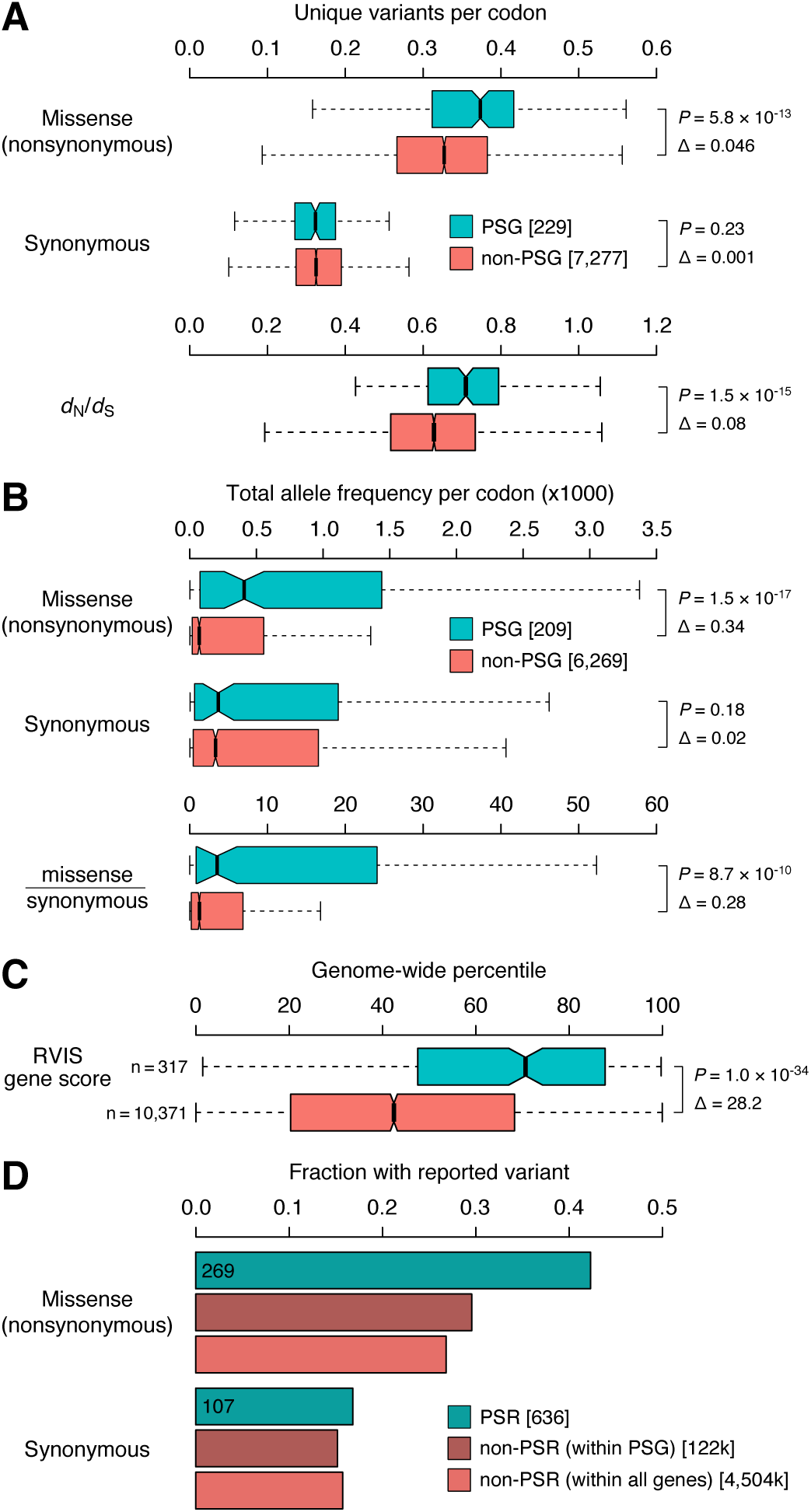
Primate positively selected genes show ongoing change in human. Genes were separated into those with strong evidence for positive selection in primates (PSG), and genes that lacked primate signatures of positive selection (non-PSG). (**A**) Distributions of the number of unique missense and synonymous variants reported per gene in ExAC (Lek et al. 2016) normalized for gene length (number of codons), as well as a *d*_N_/*d*_S_ measure of variation based on these unique variant counts (see **Methods**). (**B**) Total missense, synonymous, and missense-over-synonymous variation per gene, obtained by summing the ExAC allele frequencies of all reported variants and normalizing for gene length. (**C**) ExAC-based RVIS scores, which quantifies the amount of ‘functional variation’ (missense, nonsense and splice variants) in a gene corrected for background levels of synoynmous variation (Petrovski et al. 2013). Scores are expressed as genome-wide percentiles with higher scores indicating genes with more functional variation; i.e. a value of 71% means that 71% of genes have lower RVIS scores, thus only 29% of genes tolerate more functional variation. (**D**) Fractions of codons with variants reported in ExAC across (i) primate positively selected residues/codons (PSR) within the 209 primate PSG in our ExAC dataset, (ii) all non-PSR codons within those PSG, and (iii) all non-PSR codons, within both PSG and non-PSG. *P* values are from two-sided Mann-Whitney *U* tests; Δs represent differences between the medians.

Positively selected residues/codons (PSR) within our primate PSG show the same trends as the PSG themselves. Of the 636 PSR in genes represented in our ExAC dataset, 269 (42%) contain reported missense variants, compared to 30% for non-PSR positions in PSG genes and 27% of non-PSR positions in any gene (Figure 5D). PSR and non-PSR positions however contain similar synonymous variation (107/636 = 17% [PSR] vs. 15%/16% [non-PSR], Figure 5D). Thus, while whole-gene *d*_N_/*d*_S_ ratios are small and suggest that the majority of codons are evolving under purifying selection both between different primates and within human, the genes and codons that do evolve under positive selection in primates consistently show strong patterns of ongoing change in human populations as well.

## Discussion

In this study, we have presented an unbiased screen to identify novel virus-host genetic conflicts on the basis of statistical evidence for molecular adaptation between species. While previous investigations of virus-host genetic conflicts have focused on single genes (Sawyer et al. 2005; Elde et al. 2009; Mitchell et al. 2012; Patel et al. 2012) or subsets of genes known to be important for infection (Ortiz et al. 2009; Bozek and Lengauer 2010; Enard et al. 2016), we took a generic approach starting from entire genomes. This was enabled by the completion of several primate genome sequences bridging key evolutionary timescales between previously available species.

Earlier estimates of primate genes undergoing positive selection range from ∼1% to 4-6% to ∼10% depending on the species studied, the detection method and the number of genes tested (Bustamante et al. 2005; Chimpanzee Sequencing and Analysis Consortium 2005; Nielsen et al. 2005; Bakewell et al. 2007; Rhesus Macaque Genome Sequencing and Analysis Consortium et al. 2007; Kosiol et al. 2008; Schneider et al. 2009; George et al. 2011). Lower estimates were likely underpowered and limited by genome availability (e.g. comparing human-chimp), while higher estimates tend to be affected by the high rate of false positive predictions discussed before (Wong et al. 2008; Schneider et al. 2009; Fletcher and Yang 2010; Markova-Raina and Petrov 2011; Jordan and Goldman 2012; Privman et al. 2012; Moretti et al. 2014). Our observed *d*_N_/*d*_S_ rate across one-to-one protein-coding orthologs (0.21) is similar though slightly lower than previous estimates based on human, chimpanzee and macaque [0.23-0.26 (Chimpanzee Sequencing and Analysis Consortium 2005; Bakewell et al. 2007; Rhesus Macaque Genome Sequencing and Analysis Consortium et al. 2007)]. Given our conservative approach (minimizing false positives while perhaps missing potentially interesting sites) applied to half of all human protein-coding genes, our estimate that 3% of genes and 0.014% of codons are under positive selection should represent a reliable lower limit of what is detectable using the current whole-genome sequenced primates. Positive selection between species as a result of adaptation to a single large environmental change is typically thought to be followed by some degree of fixation of the newly acquired beneficial variants through purifying selection within populations or species (Sabeti et al. 2006; Fu and Akey 2013)(**Text S1**). Such ‘selective sweeps’ are thus expected to reduce the nonsynonymous over synonymous variation present within a single species at codons that show high *d*_N_/*d*_S_ between species. This model is assessed by for instance the McDonald-Kreitman test, which estimates how much of the variation between species is driven to fixation within species (Fay 2011). In contrast, ongoing environmental changes, such as in the case of the abundance of rapidly mutating viruses and other pathogens, will cause a recurrent selective pressure and induce adaptive changes even within a species (Daugherty and Malik 2012). Our analyses of human genetic variation demonstrate that positively selected genes and codons in primates tend to still show elevated rather than reduced levels of missense over synonymous substitutions within humans. Although it is unclear whether this indicates relaxed negative selection or persistent positive selection, these data do argue against fixation and in favor of recurrent, ongoing change in many of the primate PSG in the human population.

Genomes evolve under a large variety of selective pressures, only some of which are caused by viruses and other pathogens. Furthermore, not all viruses will have imposed sufficiently strong, long-lasting and recurrent selective pressures to leave detectable marks in the genome. Nevertheless, it is likely that our investigation of primate genomes has detected traces of genetic conflicts driven by exposure to past viruses, some of which may also influence susceptibility to present-day viruses. As most species are currently represented by a single consensus sequence, further studies of genetic variation in different human populations (our human analyses are based on global variation, independent of populations), in other primate species (de Manuel et al. 2016) and in archaic hominins (e.g. Neanderthals, Denisovans) may identify targets of positive selection caused by more recent viruses and other selective pressures (Fu and Akey 2013; Grossman et al. 2013; Mathieson et al. 2015). For example, human SNPs that correlate with current geographic virus diversity were shown to be enriched for immune genes (Fumagalli et al. 2010). Further insights into the forces that drive selection of genetic variants would require sequences from a continuum of individuals, populations and species.

Despite the strict nature of our computational procedure, our manual inspection of all positive selection hits revealed a sizeable number of artefacts. This suggests there still exists a large discrepancy between data availability and the reliability of large-scale comparative sequence analysis. Despite their limited evolutionary distance, even the study of primate genomes encounters many challenges. The large majority of positive selection artefacts remaining after stringent automated filtering arise from alignment of coding sequences that are not strictly from the orthologous genomic region in different species, i.e. they arise from alignment of non-orthologous codons. The predominant underlying source of these issues are inconsistencies in gene models and genome annotation, which causes differences in coding sequence start / stop locations, exon boundaries, pseudogene predictions and alternative transcripts. Thus, rigorous manual inspection and curation at all stages of automated pipelines remain critical for reliable results and provide insights into the current challenges in comparative genomics studies.

Our positive selection approach successfully identifies genes with a role in immunity and offer insights into other functions and cellular systems that are evolving adaptively. A striking number of positively selected genes (PSG) act as cell surface receptors in adaptive immunity. Others are involved in cytokine signaling and innate immune functions such as the complement system, intracellular pathogen recognition (both receptors and pathway members) or intrinsic antiviral activity. We also found adaptive signatures potentially related to other phenotypes that may be of interest, including morphology (hair protein *KRTAP24-1*), dietary diversity (smell and taste receptors), cholesterol and lipid transport (apolipoproteins), and energy metabolism (mitochondrial proteins).

It is peculiar to observe mitochondrial proteins among the list, as these might be regarded as having conserved housekeeping functions. One explanation is the occurrence of accelerated compensatory evolution (Osada and Akashi 2012), in which nuclear-encoded genes adapt to slightly deleterious mutations in the mitochondrial genome that are brought about by its relatively high mutation rate and the lack of recombination. Indeed, of the 13 PSG that are targeted to the mitochondrial matrix and whose direct interactors are known, 12 interact directly with either a mitochondrial rRNA, tRNA or a mitochondrial-encoded protein (**Table S10**). We further examined whether not only the protein, but also the positively selected residue interacted with a mitochondrial-encoded RNA or peptide. This appeared to be the case in two out of the five cases where structure data were available for the protein itself or for a homolog (**Table S10**).

Another exciting group of PSG are those involved in centromere structure, chromosome segregation and meiosis, which may be related to the phenomenon of centromere meiotic drive (Malik and Henikoff 2009). Meiosis in female animals is asymmetric in that only one of four haploid gametes are retained. Retention of individual chromosomes depends on a preferred binding orientation of the centromeres to the microtubular spindle via the kinetochore. Centromere evolution has been theorized to be under strong Darwinian selection because ‘selfish’ centromeres with a favored retention in meiosis are directly more likely to be transmitted (Malik and Henikoff 2009). This process may have driven the evolution of the highly variable centromere DNA sequence, marked by a specific chromatin structure containing the CENPA histone H3 variant (Malik and Henikoff 2009; Verdaasdonk and Bloom 2011). We found positive selection signatures in the centromere-associated proteins CENPO, which plays a role in linking CENPA nucleosomes to the kinetochore, and CENPT, which directly connects the centromeric DNA to the kinetochore through its histone-like fold (Nishino et al. 2012). In addition, our PSG contain two genes involved in the meiotic cohesion complex (*REC8, SGOL2*) and various genes involved in the centrosome and spindle machinery (*CEP250, PCNT, CDK5RAP2, MISP*). The rapid evolution of the centromere may be a driver contributing to the observed adaptations in these genes, even though a direct physical interaction with the centromere has not been established in all cases.

Sites under positive selection are probably important for the biological processes that led to their selection. However, deconvoluting the contributions of specific selective pressures to the complex landscape of genome variation requires additional information. We integrated the genomic signals of positive selection with orthogonal data describing the virus-host interaction to not only predict novel virus-host genetic conflicts, but also to identify positions likely important in these interactions (**Table S8**). Protein structure analyses revealed close contacts between virus particles and positively selected residues in cellular entry receptors for HIV, measles, adenoviruses and picornaviruses (*CD4, CD46, CD55*). These observations, guided by systematic analysis of virus-host evolutionary genetics, suggest positions in both viral proteins and their human receptors that are important for infection and may represent novel targets for vaccine or antibody development.

## Materials and Methods

### Sequences

We obtained protein-coding DNA sequences of all nine simian primates for which high-coverage whole-genome sequences are currently available from Ensembl release 78, December 2014 (Yates et al. 2016)(**Table S1**, **Figure S1, Supplementary Files** at http://www.cmbi.umcn.nl/∼rvdlee/positive_selection/). We processed orthology definitions from the Ensembl Compara pipeline (Vilella et al. 2009) to obtain 11,170 one-to-one ortholog clusters containing for all nine primates a single coding sequence corresponding to the canonical transcript, which usually encodes the longest translated protein (**Table S2**). Clusters consist of genes for which only one copy is found in each species, and these genes are one-to-one orthologs to the human gene. Sequences are of high quality as indicated by the general lack of undetermined nucleotides (‘N’): 98,420 of 100,530 sequences (98%) and 9,312 of 11,170 (83%) clusters contain no Ns. Genomes with most Ns are gibbon (in 680 sequences), gorilla (313) and marmoset (291). Ortholog clusters never contain sequences with internal (premature) stop codons and stop codons at the end of sequences were removed.

### Initial alignments

We first obtained multiple alignments of the clusters of orthologous primate protein sequences using MUSCLE (Edgar 2004) and Mafft (Katoh and Standley 2013), and from the Compara pipeline (i.e. filtering the larger vertebrate alignment for primate sequences)(Vilella et al. 2009). Inspections revealed that misalignment of nonhomologous codons affects virtually all alignments, as was observed in previous studies (Schneider et al. 2009; Fletcher and Yang 2010; Markova-Raina and Petrov 2011; Jordan and Goldman 2012; Privman et al. 2012). This is probably the result of the tendency of alignment algorithms to overalign, i.e. produce alignments that are shorter than the true solution due to collapsed insertions in an attempt to avoid gap penalties (Löytynoja and Goldman 2008; Fletcher and Yang 2010). The PRANK algorithm to some extent prevents alignment of nonhomologous regions by flagging gaps made during different stages of progressive alignment and permitting their reuse without further penalties (Löytynoja and Goldman 2008). As PRANK has been shown to provide better initial alignments for large-scale positive selection detection (Schneider et al. 2009; Fletcher and Yang 2010; Markova-Raina and Petrov 2011; Jordan and Goldman 2012; Privman et al. 2012; Moretti et al. 2014), we obtained multiple alignments of the primate ortholog clusters using the PRANK codon mode (`prank +F –codon`; v.140603). We used the default settings of (i) obtaining a guide tree from MAFFT for the progressive alignment procedure and (ii) selecting the best alignment from five iterations. These settings likely result in the best alignment for a given cluster of sequences (including those showing a gene tree topology that differs from the species tree topology). The PRANK approach markedly improved the initial alignments.

### Alignment filtering and masking

Even with improved initial alignments, positive selection studies remain affected by a high rate of false positive predictions. Part of those may be alleviated by additional automated masking of unreliable alignment regions. GUIDANCE assesses the sensitivity of the alignment to perturbations of the guide tree (Penn et al. 2010) and has been recommended for positive selection studies (Jordan and Goldman 2012; Privman et al. 2012; Moretti et al. 2014). We applied GUIDANCE with the default 100 bootstrap tree iterations (`guidance.pl --program GUIDANCE --seqType nuc --msaProgram PRANK --MSA_Param “\+F \-codon”`; v1.5). TCS assesses alignment stability by independently re-aligning all possible pairs of sequences and scoring positions through comparison with the multiple alignment (Chang et al. 2014). We ran TCS on translated PRANK codon alignments (`t_coffee -other_pg seq_reformat -action +translate`; `t_coffee -evaluate -method proba_pair -output score_ascii, score_html`; Version_11.00.61eb9e4).

Low confidence scores of either method led us to remove entire alignments from our analysis or mask individual columns and codons. Alignments were removed in the case of a low score (default cutoffs of <60% for GUIDANCE, <50% for TCS) for (i) the overall alignment or (ii) one or more sequences (i.e. we only retained alignments with sequences for all nine species). Entire columns were masked if GUIDANCE <93% or TCS <4; individual codons were masked if <90% or <4. Masked nucleotides were converted to ‘n’ characters to distinguish them from undetermined nucleotides in the genome assemblies (‘N’). For visualization and quality inspection purposes we translated the masked codon alignments to the corresponding protein alignment. Nucleotides ‘n’ and ‘N’ were converted to ‘o’ and ‘X’ upon translation, respectively. Detailed visual inspection revealed the value of our masking approach: masked codons tend to comprise unreliable alignment regions, primarily consisting of large inserts, insertion-deletion boundaries (i.e. regions bordering well-aligned blocks), and aligned but nonhomologous codons (**Supplementary Files**).

### Evolutionary analyses: reference phylogenetic tree

Maximum likelihood (ML) *d*_N_/*d*_S_ analysis to infer positive selection of genes and codons was performed with codeml of the PAML software package v4.8a (Yang 2007)(**Text S1**). We used a single phylogenetic tree with branch lengths for the ML analysis of all alignments to limit the influence of gene-specific phylogenetic variability. To obtain this reference tree, we concatenated all 11,096 masked alignments into one large alignment and ran the codeml M0 model (i.e. fitting a single *d*_N_/*d*_S_ for all sites; `NSsites = 0, model = 0, method = 1, fix_blength = 0`), provided with the well-supported topology of the primate phylogeny (Perelman et al. 2011; Yates et al. 2016). We took this approach for two main reasons: (i) to best reflect the overall evolutionary distance between the primate species (which influences codon transition probabilities in the ML calculations, **Text S1**), and (ii) to estimate branch lengths in units compatible with codon-based evolutionary analyses, i.e. the number of nucleotide substitutions per codon. For comparisons with other primate phylogenetic trees, the branch lengths of our codon-based tree were converted to nucleotide substitutions per site (i.e. nucleotide substitutions per codon divided by three). The codeml M0 model under the F61 or F3X4 codon frequency parameters resulted in virtually identical phylogenetic trees (median branch length difference of a factor 0.99) and *d*_N_/*d*_S_ estimates (0.213 vs. 0.217; **Figure S1, Supplementary Files**). The M0 tree is also highly similar to a ML phylogenetic tree inferred from the same concatenated alignment using nucleotide rather than codon substitution evolutionary models (median branch length difference of a factor 0.98; **Figure S1**; RAxML v7.2.8a (Stamatakis 2014); `-f a -m GTRCAT -N 100`).

### Evolutionary analyses: inference of positive selection

In the first of two steps for inferring positive selection using codeml, the 11,096 filtered and masked alignments were subjected to ML analysis under evolutionary models that limit *d*_N_/*d*_S_ to range from 0 to 1 (‘neutral’ model) and under models that allow *d*_N_/*d*_S_ > 1 (‘selection’ model; **Text S1**)(Yang and Bielawski 2000). Genes were inferred to have evolved under positive selection if the likelihood ratio test (LRT) indicates that the selection model provides a significantly better fit to the data than does the neutral model (*P*_LRT_ < 0.05, after Benjamini Hochberg correction for testing 11,096 genes). We included apparent Positively Selected Genes (aPSG) if they met the LRT significance criteria under all four tested ML parameter combinations. These combinations consist of two sets of evolutionary models: M1a (neutral) vs. M2a (selection); M7 (beta) vs. M8 (beta&ω). And two codon frequency models: F61 (empirical estimates for the frequency of each codon); F3X4 (calculated from the average nucleotide frequencies at the three codon positions). I.e. we used combinations of the following codeml parameters: `NSsites = 1 2` or `NSsites = 7 8`; `CodonFreq = 2` or `CodonFreq = 3`; `cleandata = 0, method = 0, fix_blength = 2`. 2,992 (27%) genes showed significant evidence of apparent positive selection at the level of the whole alignment (**Figure S2A**).

Second, for the significant aPSG we retrieved from the site-specific codeml ML analyses (step one, above) the Bayesian posterior probabilities, which indicate the individual codons that may have evolved under positive selection (**Text S1**)(Yang et al. 2005). We included apparent Positively Selected Residues (aPSR) if their codons were assigned high posteriors under all four ML parameter combinations (*P*_posterior_ (ω > 1) > 0.99). 416 aPSG contain at least one significant aPSR (1405 in total; **Figure S2B**).

### Quality control

We subjected each inferred aPSR and aPSG to visual inspection (**Table S3**). In this way we identified several indicators for positive selection artefacts that we then used for their automated detection in the complete set. First, we obtained the gene trees for our individual masked alignments using RAxML (Stamatakis 2014)(`-f a -m GTRGAMMAI -N 100`). Type-I [orthology] and -II [transcript definitions] artefacts tend to lead to gene trees with (i) a long-branched clade consisting of the set of sequences that are distinct from the others (e.g. paralogs, alternative exons), and (ii) a topology that is not congruent with the well-supported species phylogeny (**Figure S3**). We filtered out likely false positives by selecting gene trees with an extreme longest/average branch length ratio. Second, to assess the distribution of PSR across exons, we mapped Ensembl exon coordinates for human transcripts to the human protein sequences. Type-II [transcript definitions] and -III [termini] artefacts could often be filtered out by a high concentration of aPSR located to a single exon (**Supplementary Files**).

### GC-biased gene conversion (gBGC)

The effects of gBGC seem specifically correlated to regions of high meiotic recombination in males rather than females (Ratnakumar et al. 2010). We calculated genomic overlaps of PSG and non-PSG with male (8.2% of PSG, 7.7% of non-PSG) and female (6.7% of PSG, 8.1% of non-PSG) recombination hotspots in human, which we obtained from the family-based deCODE maps (Kong et al. 2010) via the UCSC genome browser (Rosenbloom et al. 2015). Sex-averaged recombination hotspots estimated from linkage disequilibrium patterns were obtained from HapMap Release 22 (International HapMap Consortium et al. 2007)(43% of PSG, 39% of non-PSG). Human genomic regions under the influence of gBGC were predicted by phastBias (Capra et al. 2013)(9.1% of PSG, 11.4% of non-PSG).

### PSG function analyses

Our final curated set of 331 PSG (**Table S4**, **S8**) was analyzed for gene ontology terms, pathways databases and other functions (**Table S6**, **S7**) (Huang et al. 2009; Alonso et al. 2015), compared to a background of 11,011 genes (the 11,096 genes tested for positive selection excluding the 85 artefacts). Enrichment statistics were calculated for overlaps between the PSG and various gene lists (**Table S8**), by assessing whether a gene list of interest was significantly over-represented among the 331 PSG compared to the 10,680 non-PSG (two-tailed Fisher’s exact test on a 2x2 contingency table of the overlap). Secreted (keyword KW-0964) and membrane (KW-0472) proteins were obtained from UniProt (UniProt Consortium 2014). Innate immunity, PRR and virus-human PPI gene lists are described in (van der Lee et al. 2015). We mined the GenomeRNAi database (Schmidt et al. 2013) for genes whose perturbation significantly affects viral infection or replication. Data on gene expression in mouse immune cell subsets were obtained from the Immunological Genome Project (using the recommended expression thresholds – usually 120)(Shay and Kang 2013) and mapped to human orthologs. Optimized atomic-resolution protein structures of virus-host interactions were obtained from PDB_REDO (Joosten et al. 2009), visualized with YASARA (www.yasara.org), and analyzed using WHAT IF (Vriend 1990).

### Human variation

To study genetic variation in the human population we obtained all variants and allele frequencies reported by the Exome Aggregation Consortium (ExAC, release 0.3.1) (Lek et al. 2016) from the variant call file (VCF). ExAC combined over 91,000 unaffected and unrelated exomes from a range of primarily disease-focused consortia, into 60,706 high-quality exomes. These exomes originate from diverse populations, including European (Non-Finnish and Finnish), African, South Asian, East Asian, Latino. Our analyses focus on global human variation rather than stratified populations, but it should be noted that African populations have greater levels of genetic diversity and that Middle Eastern and Central Asian samples are underrepresented in ExAC (Lek et al. 2016). ExAC variants were mapped from genomic position to codons using an in-house pipeline that matches the longest translated GENCODE (Harrow et al. 2012) sequence for each mRNA-validated protein-coding gene to a Swiss-Prot (UniProt Consortium 2015) protein sequence. This ExAC dataset was then merged with the total set of genes tested for positive selection in our primate analysis. The final dataset (**Table S11**) covers 7,506 genes of which 229 are primate PSG (containing 636 PSR).

We considered only single nucleotide variants (SNVs) and included all SNVs that pass the filters imposed by ExAC. We did not impose any allele frequency filters and used the aggregated allele counts/frequencies across all populations. (i) At the level of unique variants, we first calculated the number of unique missense and synonymous variants reported per codon by aggregating the potentially multiple, distinct variants reported at the same codon. We then summed these per-codon counts to obtain the number of unique variants per gene, which we normalized for gene length by dividing by the number of codons (“Unique variants per codon”, Figure 5A). A *d*_N_/*d*_S_ measure of human variation was obtained by correcting the observed unique missense and synonymous variant counts in a gene (observed, obs) for the total possible missense and synonymous variants in that gene based on the codon table (background, bg): 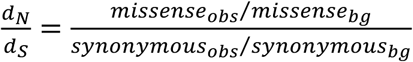. Note that this ExAC-based human *d*_N_ /*d*_S_ uses a background of possible variants based exclusively on the codon table, ignoring any biases that influence codon transition probabilities, such as transition/transversion rates, codon frequencies and divergences times, which are all taken into account in our analyses of primate positive selection using codeml. These differences in approach may partly explain the overall higher *d*_N_/*d*_S_ observed in our analyses of human compared to primates. (ii) At the level of allele frequencies (AF), we measured total human variation in a gene by summing the reported AFs (counts for the alternative allele / allele number [total number of called genotypes]), separately for missense and synonymous variants. These aggregated AFs were normalized for gene length to obtain the average (per-codon) allele frequency of all missense and all synonymous variation within the gene (“Total allele frequency per codon”, Figure 5B).

ExAC-based Residual Variation Intolerance Scores (RVIS) were downloaded from http://genic-intolerance.org/data/GenicIntolerance_v3_12Mar16.txt. RVIS is a regression-based measurement of the tolerance for a gene to accumulate ‘functional variation’ (missense, nonsense and splice variants), given its level of synonymous variation (Petrovski et al. 2013). We analyzed the ‘default’ RVIS genome-wide percentiles, which were calculated based on variants with a MAF of at least 0.05% in at least one of the six individual ethnic strata from ExAC (i.e. we used *%ExAC_0.05%popn*), but all MAF filters gave similar results.

### Scripts and tools

Our procedures and analyses consists of custom Perl and R scripts, available upon request. In addition to the methods cited above, we made extensive use of the Ensembl API version 78 (Yates et al. 2016), GNU Parallel (Tange 2011) and Jalview (Waterhouse et al. 2009).

## Acknowledgments

We would like to thank Dei M. Elurbe, Manja Leemans, Colin Logie, Eelco Tromer, Geert Kops and Berend Snel for valuable discussions. This work was supported by the Virgo consortium, funded by the Dutch government (FES0908), and by the Netherlands Genomics Initiative (050-060-452).

